# Rapid evolution of promoters from germline-specifically expressed genes including transposon silencing factors

**DOI:** 10.1101/2023.11.01.564449

**Authors:** David W. J. McQuarrie, Azad Alizada, Benjamin Czech Nicholson, Matthias Soller

## Abstract

**Background:** The piRNA pathway in animal gonads functions as an ‘RNA-based immune system’, serving to silence transposable elements and prevent inheritance of novel invaders. In *Drosophila*, this pathway relies on three gonad-specific Argonaute proteins (Argonaute-3, Aubergine and Piwi) that associate with 23-28 nucleotide piRNAs, directing the silencing of transposon-derived transcripts. Transposons constitute a primary driver of genome evolution, yet the evolution of piRNA pathway factors has not received in-depth exploration. Specifically, channel nuclear pore proteins, which impact piRNA processing, exhibit regions of rapid evolution in their promoters. Consequently, the question arises whether such a mode of evolution is a general feature of transposon silencing pathways.

**Results:** By employing genomic analysis of coding and promoter regions within genes that function in transposon silencing in *Drosophila*, we demonstrate that the promoters of germ cell-specific piRNA factors are undergoing rapid evolution. Our findings indicate that rapid promoter evolution is a common trait among piRNA factors engaged in germline silencing across insect species, potentially contributing to gene expression divergence in closely related taxa. Furthermore, we observe that the promoters of genes exclusively expressed in germ cells generally exhibit rapid evolution, with some divergence in gene expression.

**Conclusion:** Our results suggest that increased germline promoter evolution, in partnership with other factors, could contribute to transposon silencing and evolution of species through differential expression of genes driven by invading transposons.

## Background

Eukaryotic organisms are continuously challenged from genomic parasites called transposable elements (TEs) [1–4]. Unchecked transposon activity often results in reduced reproductive fitness [5–8]. To negate the detrimental effects posed by TE mobilisation, vital regulatory pathways evolved that efficiently suppress transposon activity. At the centre of these are 23-28 nucleotide small RNAs, called PIWI-interacting RNAs (piRNAs). piRNAs are generated from transposons and from genomic clusters and bind to Argonaute proteins of the PIWI-clade, including P-element induced wimpy testis (Piwi). Aubergine (Aub) and Argonaute-3 (Ago3). The piRNA pathway is often referred to as an ‘RNA-based immune system’ thanks to its ability to adopt to new TE insertions and the memory of past transposon activity through piRNA clusters through RNA transgenerational inheritance [1, 2, 9, 10].

The piRNA pathway is arguably best understood in the ovary of *Drosophila melanogaster*. It was shown that ovaries feature two branches of the silencing machinery [5], one that is active in germ cells and a condensed version that is specific to somatic follicle cells that surround the germline [2, 11]. Somatic follicle cells only feature unistrand piRNA clusters, such as *flamenco* (*flam*), and exclusively produce piRNAs via a Zucchini (Zuc)-dependent biogenesis mechanism that takes place on the mitochondria surface involving Armitage (Armi), Papi, Vreteno (Vret), Gasz, Daedalus (Daed), Minotaur (Mino), Shutdown (Shu), and Sister of Yb (SoYb) (Fig. 1A) [3, 12, 13]. Licensing of *flam* for piRNA production takes place upstream, at so-called Yb-bodies [1, 12]. Somatic cells only express Piwi, which following the association with a mature piRNA shuttles to the nucleus. There, Piwi-piRNA complexes scan for nascent TE transcripts and instruct co-transcriptional gene silencing (cTGS) that requires components of the general chromatin silencing machinery and the piRNA pathway-specific factors Asterix (Arx), Piwi, Panoramix (Panx), Nuclear export factor 2 (Nxf2) and Maelstrom (Mael) (Fig. 1A) [1, 2, 12, 14].

**Fig. 1:**
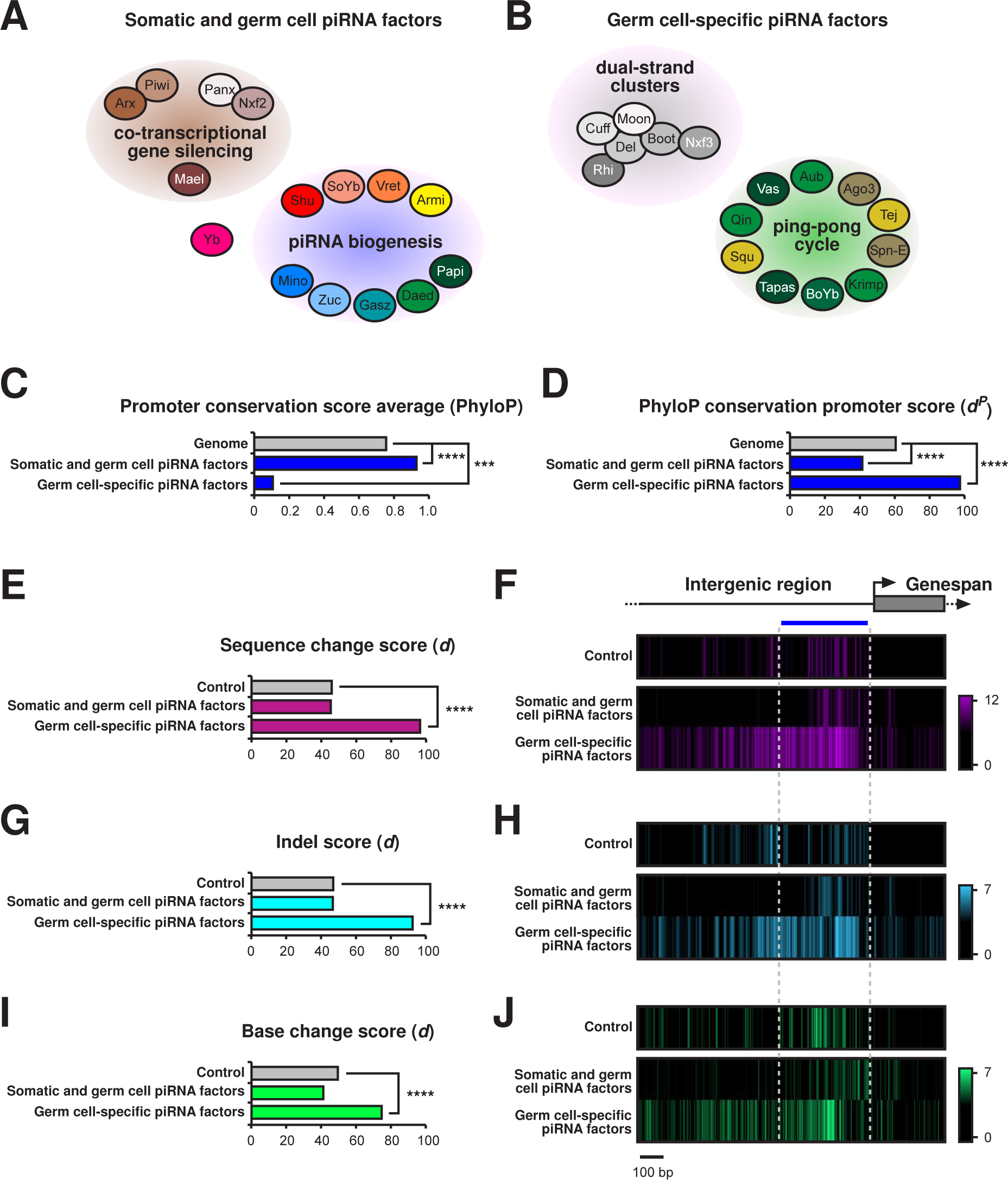
Promoter regions of germ cell-expressed piRNA processing factors are hot spots for rapid evolution. A and B) Schematic depiction of somatic and germ cell piRNA factors involved in co-transcriptional gene silencing and piRNA biogenesis (A) and germline-specific piRNA pathway genes involved in dual-strand cluster regulation and the ping-pong cycle (B) in the *Drosophila* ovary. C and D) PhyloP27way conservation score averages (C) and PhyloP27way conservation promoter *d^P^* scores (D) for the 350 nucleotide promoter regions of the somatic and germ cell piRNA factors and the germ cell-specific piRNA factors compared to all genes in the *Drosophila* genome. Statistically significant differences from unpaired student t-tests (C) and non-parametric chi-squared tests (D) are indicated by asterisks (*** p≤0.001, **** p≤0.0001 following Bonferroni correction). E-J) Heatmaps indicating sequence (F, purple), indel (H, blue) or base change (J, green) accumulation, and their quantification (E-I) depicted as *d* scores for each of the analysed gene groups compared to the control among closely related *D. melanogaster*, *D. simulans*, *D. sechellia*, *D. yakuba* and *D. erecta*. Regions of 1000 nucleotides upstream and 300 nucleotides downstream of the TSS were analysed. Analysis was performed for the control group (m^6^A writer complex and readers), the somatic and germ cell piRNA factors, and the germ cell-specific piRNA factors. The blue line indicates the promoter region used for quantification of the substitution rate based on the gene and intergenic region schematic. Statistically significant differences from non-parametric chi-squared tests are indicated by asterisks (**** p≤0.0001 following Bonferroni correction).

In germ cells, piRNAs predominantly originate from dual-strand clusters and are produced by a different biogenesis mechanism called the ping-pong amplification cycle (Fig. 1B) [1–3, 5, 11, 12, 15]. Dual-strand clusters harbour remnants of transposons and are expressed via a non-canonical transcription process that is centred on a complex containing the HP1a homolog Rhino (Rhi), Deadlock (Del) and Cutoff (Cuff), and further requires Moonshiner (Moon) (Fig. 1B). This Rhi-Del-Cuff complex also recruits a non-canonical export machinery consisting of Bootlegger (Boot) and Nuclear export factor 3 (Nxf3), which transport piRNA precursors for processing to a germ-cell specific perinuclear region termed “nuage” (Fig. 1B). Here, the ping-pong cycle takes place and relies on the two PIWI proteins Aubergine (Aub) and Argonaute 3 (Ago3), as well as several RNA binding proteins and Tudor-domain containing factors including Tejas (Tej), Vasa (Vas), Qin, Squash (Squ), Tapas, Brother of Yb (BoYb), Krimper (Krimp), and Spindle-E (Spn-E) (Fig. 1B) [1–3, 12, 15, 16].

Channel nuclear pore proteins (Nups) have been implicated in piRNA regulated transposon silencing in *C. elegans* [17]. Interestingly, in *Drosophila*, Nups were shown to affect transposon silencing in both the germline [16, 18], and in somatic cells of the ovary, here specifically by contributing to the conversion of precursor RNAs from the *flam* locus into piRNAs [19]. Recently, we found that inner and outer ring Nups in the nuclear pore complex (NPC) evolve rapidly through indel accumulation in their promoters [20, 21]. Such promoter indel variability in Nup54 has dominant, pleiotropic effects on sexual differentiation including neuronal wiring important for female post-mating behaviours directed by male-derived sex peptide, that as a result of sexual conflict could drive speciation by such mechanism [17, 20, 22]. Intriguingly, nuclear import/export pathways have been linked to speciation through hybrid incompatibility, but are in addition important for many cellular processes beyond piRNA processing [23, 24]. Hence, promoter evolution particularly of piRNA processing genes could have wide impact on transposon silencing and evolution of species through differential expression of genes, but whether regulators of germline transposon silencing generally evolve rapidly through their promoters has not been determined.

Here, we analyse the promoters of genes involved in the ovarian piRNA pathway. We compare piRNA factors required in somatic cells (Yb) or in both the somatic and germline compartments (cTGS and piRNA biogenesis, Fig. 1A) with those that are germ cell-specific (dual-strand clusters and the ping-pong cycle, Fig. 1B). We find that the promoters of germline-specific piRNA factors are hot spots for rapid evolution compared to those of somatic and germ cell piRNA pathway genes and the *Drosophila* genome average throughout insect species. Analysis of genes with significant accumulation of promoter changes reveals a mix of indel and base change accumulation. Further, our analysis reveals that rapid promoter evolution is a general feature of genes specifically expressed in germ cells, while soma-expressed genes evolve at a similar rate to the average of all *Drosophila* genes. Through cross-species differential ovary RNA-seq analysis we reveal that germ cell-specific genes minimally diverge in gene expression levels compared to genes expressed in somatic cells. Our findings highlight that promoters of germline genes involved in transposon silencing evolve rapidly and are accompanied by diverging gene expression, suggesting a possible mode of rapid speciation through accumulation of changes in promoters in correlation with additional regulatory factor divergence.

## Results

### Promoter regions of germline-specific piRNA factors are hot spots for rapid evolution

Since Nups have been shown to function in transposon silencing in the germline (12 from 14 tested are above the control) and display fast evolving promoters [16, 20, 21], we analysed the rate of evolution in promoters and coding regions of piRNA pathway genes. We separated all piRNA pathway genes based on their genetic requirement into two categories: piRNA factors essential in somatic cells, including the soma-specific piRNA factor Yb and piRNA factors expressed in both somatic and germline cells (cTGS and piRNA biogenesis, Fig. 1A), and germ cell-specific piRNA proteins (dual-strand clusters and ping-pong cycle, Fig 1b). The piRNA factors essential in somatic cells include *armi*, *papi*, *vret*, *Gasz*, *daed*, *mino*, *zuc*, *shu*, *SoYb*, *Yb*, *arx*, *piwi*, *panx*, *nxf2*, and *mael* (Fig. 1A). The group of piRNA factors that are exclusively required in the germline are *aub*, *tej*, *Ago3*, *vas*, *qin*, *squ*, *tapas*, *BoYb*, *krimp*, *spn-E, Boot*, *moon*, *nxf3*, *cuff*, *rhi*, and *del* (Fig. 1B) [3, 14, 25].

We defined the promoter containing region as a 350-nucleotide (-30 to -380) window upstream of the predicted TATA box region 30 nucleotides upstream of the transcription start site (TSS) [26]. In a first approach, we performed a genome-wide promoter evolution analysis of germline-specific as well as somatic and germ cell piRNA factors and compared it to all genes in the *Drosophila* genome (Fig 1C-D). We used publicly available PhyloP27way data a conservation score. We observed a significant accumulation of changes in the germ cell-specific piRNA factor promoter regions compared to the genome (Fig. 1C-F). Of note, the somatic and germ cell piRNA factors evolved slower compared to the genome.

We next compared indel and base changes in the promoter regions between five closely related *Drosophila* species (*D. melanogaster*, *D. simulans*, *D. sechellia*, *D. yakuba* and *D. erecta*). Promoter scores (*d*) were calculated and compared to a control group comprised of the m^6^A methylation machinery genes (*Mettl3*, *Mettl14*, *fl(2)d*, *virilizer*, *flacc*, *nito* and *Hakai*, *Ythdc1* and *Ythdf*), chosen due to their high evolutionary conservation and requirement for strict stoichiometry for functionality, making this an ideal control group to monitor promoter evolution [21, 27, 28]. In this analysis, the group of somatic and germ cell piRNA factors showed similar evolution rates compared to the control (Fig. 1E-J), while germ cell-specific piRNA factors as a group evolved at a much higher rate (Fig. 1G and H), accruing a significantly increased number of promoter-located indels (Fig. 1G and H) and base changes (Fig. 1I and J). At the individual gene level, 7 out of 14 of germ cell-specific piRNA factors (*Ago3* was omitted due to its low conservation between closely related *Drosophila* species [29–31] showed significant accumulation of promoter changes (Supplementary Fig. 1A). Among the somatic and germ cell piRNA factors, 3 out of 15 genes had significant accumulation of promoter changes (Supplementary Fig. 1A).

To assess general evolution in the coding regions, we used McDonald-Kreitman tests (MKTs) and analysed polymorphisms and divergence within *D. melanogaster* and between *D. melanogaster* and *D. simulans* for two ancestral populations (Congo and Zambia) [32]. Analysis of grouped germ cell-specific piRNA factors flagged these factors as under positive selection for both populations. Conversely, the somatic and germ cell piRNA factors were not flagged as under positive selection, (Supplementary Fig. 1B).

### Promoters of genes required for germline transposon silencing are fast evolving

Given the rapid evolution of germ cell-specific piRNA factors among closely related *Drosophila* species, we wanted to determine (1) whether this is a general feature of insect evolution, and (2) whether this is a general feature of genes required for transposon silencing with pleiotropic roles that have been identified in a genetic screen (GTS100 genes), since this provides a larger gene group [16]. To get a scale of the persistence of evolutionary changes in promoters of GTS100 genes we used publicly available PhyloP27way data from UCSC genome browser [33, 34]. Genes were ordered based on transposon de-repression scores [16] and *d^P^* was calculated for individual genes, the genome average as a control group, and for the average of GTS100 gene promoter regions (Fig. 2A-C, 3A). Genes without full PhyloP coverage were removed from the analysis (where ≥1 nucleotide of the analysed genomic region contained the *D. melanogaster* sequence only).

**Fig. 2:**
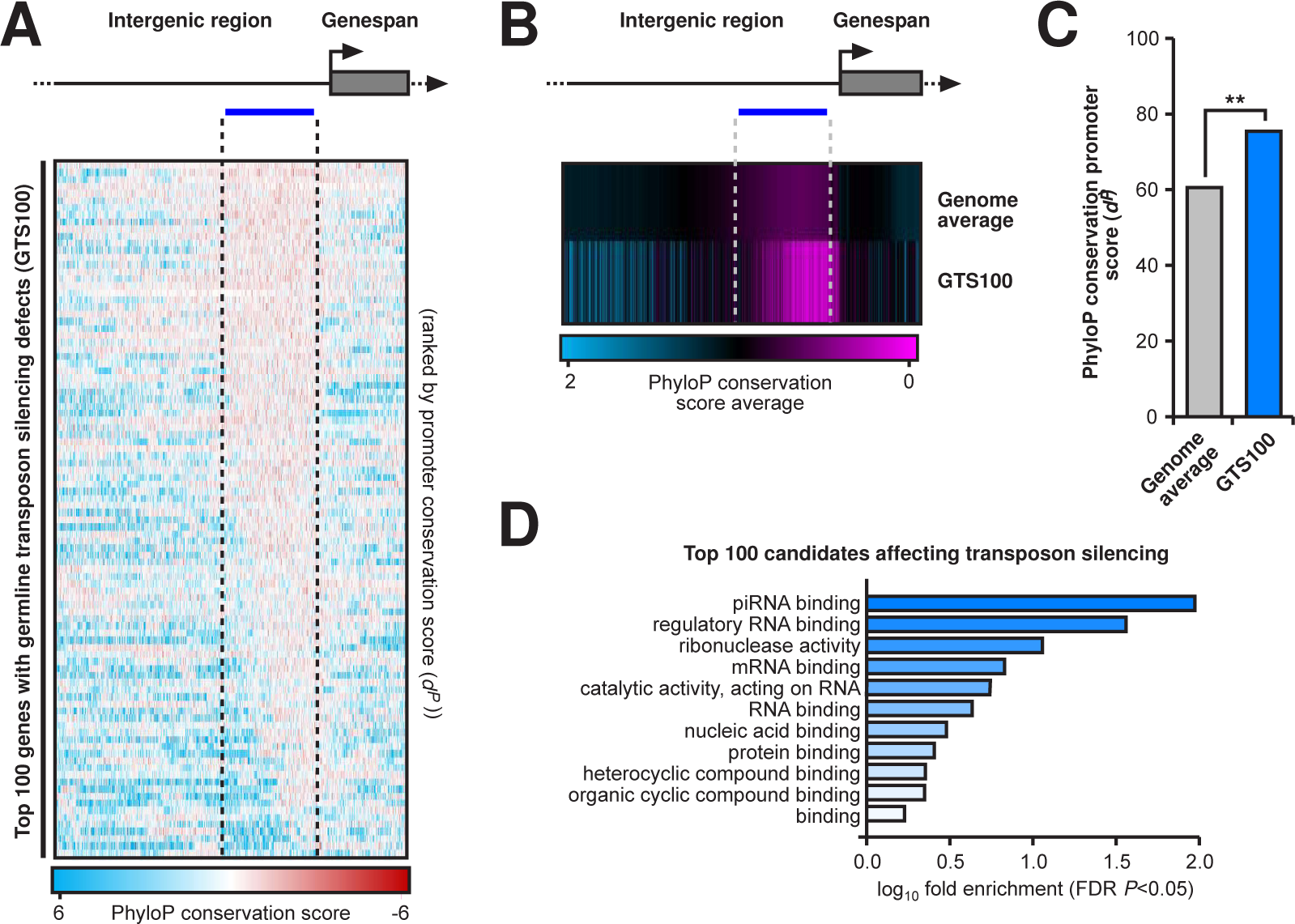
Promoters of genes required for germline transposon silencing evolve fast. A) PhyloP27way nucleotide scores for the top 100 genes affecting transposon silencing in germ cells based on transposon derepression *z* scores and ordered by promoter conservation *d^P^* scores. A region of 1000 nucleotides upstream and 300 nucleotides downstream are shown. Red represents lower conservation while blue represents higher. B) Average PhyloP27way nucleotide scores for the top 100 genes affecting transposon silencing in germ cells compared to the genome average. C) Comparison of PhyloP27way conservation promoter scores (*d^P^*) for each of the 100 genes affecting transposon silencing in germ cells compared to the genome average. Statistically significant differences from non-parametric chi-squared tests are indicated by asterisks (*** p≤0.001 following Bonferroni correction). D) Representative GO analysis for the top 100 gene candidates affecting transposon silencing.

The promoter score average of GTS100 genes showed a significant increase in promoter nucleotide changes compared to the control (Fig. 2C). Although many of these genes have additional roles, gene ontology (GO) analysis confirmed a primary role in piRNA binding (Fig. 2D) [35–37].

To understand whether GTS100 genes were individually fast evolving, individual *d^P^* scores were calculated (Fig. 3A). Compared to the genome average, 49 genes (FEPG, Fast Evolving Promoter Germline genes) were significantly increased in promoter change events, while 12 were significantly decreased (Fig. 3A). Of note *Brother of Yb* (*BoYb*) which is considered the germline replacement of *Yb* [3], also displayed a fast evolving promoter in contrast to its somatic counterpart *Yb* (Fig. 3A). To specifically look at indels at a high confidence level, we analysed FEPG genes for rapid promoter evolution between the previously used five *Drosophila* species (Fig. 3B-D). Here, indels and base changes were measured, showing a significant increase in both indels and base changes compared to the control (Fig. 3C and D).

**Fig. 3:**
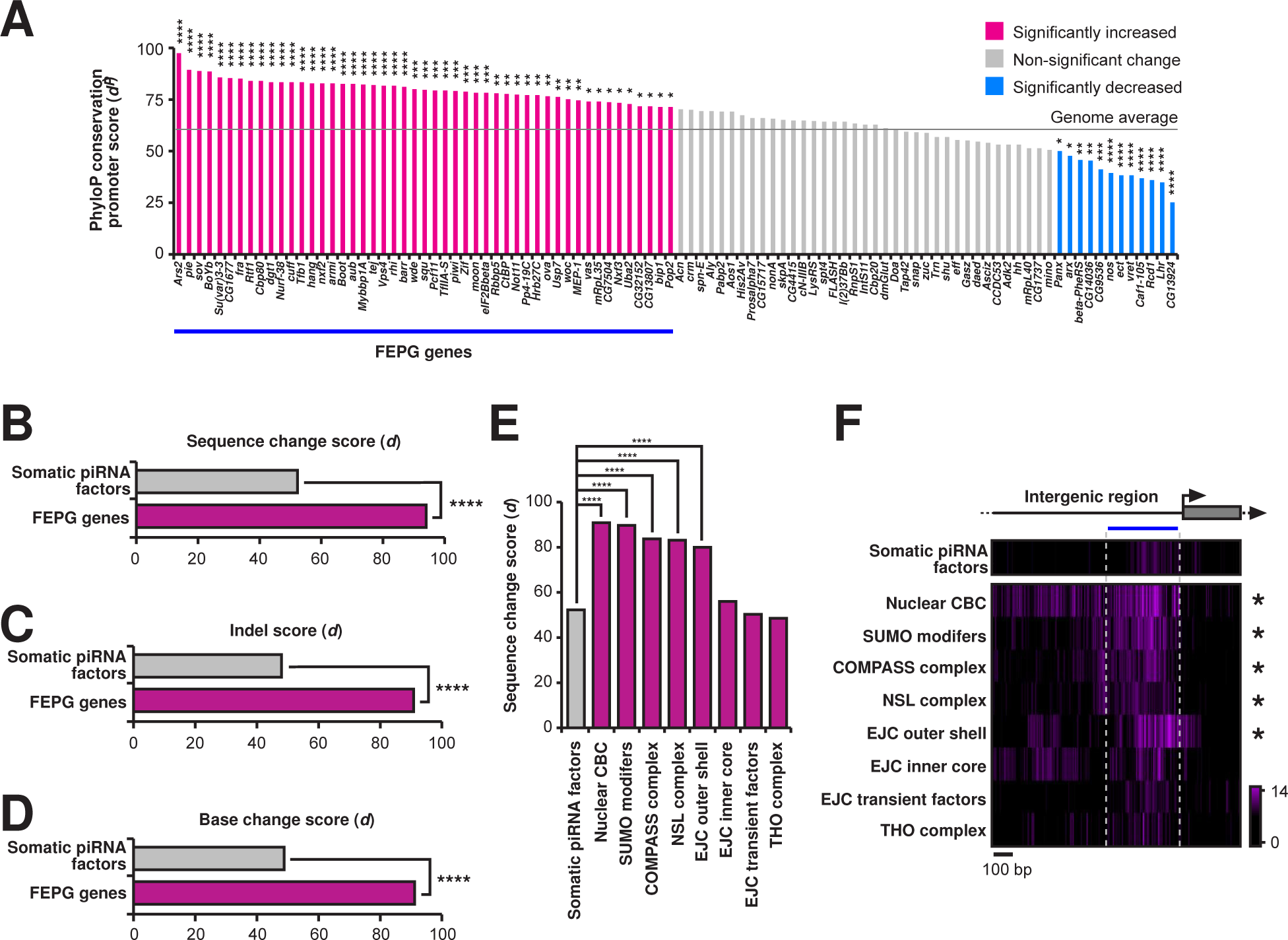
Promoters of FEPG genes and their associated complexes are hotspots for rapid evolution. A) Individual gene comparison of PhyloP27way conservation promoter scores (*d^P^*) for the top 100 genes affecting transposon silencing in germ cells compared to the genome average. Statistically significant differences from non-parametric chi-squared tests are indicated by asterisks (* p≤0.05, ** p≤0.01, *** p≤0.001, **** p≤0.0001 following Bonferroni correction). B-D) Sequence analysis for change accumulation among closely related *D. melanogaster*, *D. simulans*, *D. sechellia*, *D. yakuba* and *D. erecta*. Comparisons were performed for sequence change (B), indel (C), and base change (D) *d* scores for each of the positive 49 genes affecting transposon silencing in FEPG gene group compared to the somatic and germ cell piRNA factors. Statistically significant differences from non-parametric chi-squared tests are indicated by asterisks (**** p≤0.0001 following Bonferroni correction). E) Comparison of *d* scores for protein complex genes involved in germ cell transposon silencing compared to the somatic and germ cell piRNA factors. Statistically significant differences from non-parametric chi-squared tests are indicated by asterisks (**** p≤0.0001 following Bonferroni correction). F) Heatmaps indicating sequence change accumulation for protein complex genes involved in germ cell transposon silencing compared to the piRNA factors shown in purple among closely related *D. melanogaster*, *D. simulans*, *D. sechellia*, *D. yakuba* and *D. erecta*. Regions of 1000 nucleotides upstream and downstream of the TSS were analysed. The blue line indicates the promoter region used for quantification of the substitution rate.

### A subset of piRNA factors that are part of distinct protein complexes display hot spots for rapid evolution in promoters

In a saturating genetic screen for regulators of transposon silencing in germ cells members of the Nup complex were identified as regulators of germline piRNA silencing, and later shown to rapidly evolve through their promoters [16, 20, 21]. In the same screen, numerous genes whose products are part of distinct protein complexes were identified, including the nuclear cap-binding complex (CBC), THO complex, the exon junction complex (EJC) inner core, outer shell and EJC transient factors, the non-specific lethal (NSL) complex, SUMOylation modifiers and the complex of proteins associated with Set1 (COMPASS) (Supplementary Fig. 2). Given the fast evolution of promoters in germline-specific piRNA pathway genes and the Nup complex, we analysed promoters of the additional protein complexes affecting germline piRNA silencing to see whether their rate of evolution matched that of other piRNA silencing factors. Compared to somatic and germ cell piRNA factors, promoter regions for the nuclear CBC, SUMO modifiers, COMPASS complex, NSL complex and EJC outer shell showed a significant accumulation of sequence changes (Fig. 3E and F). Analysis of individual genes from both significant and non-significant gene groups revealed that 43% of analysed genes have rapidly accumulated sequence changes in their promoters (Supplementary Fig. 2). Specifically looking at genes in fast evolving complexes (Fig. 3E and F) revealed that 56% of genes in these groups rapidly evolve through accumulation of mutations in their promoters (Supplementary Fig. 2).

When analysing the base changes and indels of rapid evolving promoters (Supplementary Fig. 3A-D), indel accumulation was observed for the SUMO modifiers, COMPASS complex, NSL complex and EJC outer shell (4 out of 5), while the nuclear CBC showed a significant decrease (Supplementary Fig. 3B). The nuclear CBC, SUMO modifiers and COMPASS complex also showed a significant accumulation of base changes in their promoter regions (3 out of 5) (Supplementary Fig. 3D). Individually, 4 out of 27 genes were significantly enriched in indel events (*Hcf*, *dgt1*, *nsl1*, and *Bin1*), while base change accumulation events indicated 8 out of 27 genes (*Ars2*, *cbp80*, *Ulp1*, *Set1*, *Hcf*, *Cfp1*, *MBD-R2*) being significantly increased (Supplementary Fig. 3E).

Next, we analysed the rate of evolution in the coding regions of these complexes using MKT tests [32]. This analysis flagged the EJC outer shell, EJC transient facts, NSL complex, and SUMO modifiers as under positive selection in both populations, while the Nuclear CBC was under positive selection in the Zambia population only (Supplementary Fig. 4).

### Analysis of piRNA factor differential gene expression in ovaries of *D. melanogaster* and *D. yakuba*

Since promoters of FEPG genes are fast evolving, we analysed publicly available RNA-seq data from ovaries between *D. melanogaster* and *D. yakuba* to assess gene expression divergence in the FEPG gene group compared to the somatic and germ cell piRNA factors (Fig. 4A) [38–40]. Here, many FEPG genes had a tendency for slight significant divergence in gene expression in both directions (*D. melanogaster* or *D. yakuba*), but this was true for only few somatic and germ cell piRNA factors genes including *arx* (Fig. 4A). Notably, we observed large expression divergence for *CG32152, MEP-1* and *Rbbp5* (Fig. 4A). Aligning the promoters of these three genes revealed an increased number of indels and base changes compared to the 5’UTR or the coding region (Fig. 4B-D), but the cohort was unfortunately too small to detect whether increased changes in promoters correlate with altered expression between the two species. Further limitations stem from the pleiotropic roles of the analysed genes, and therefore expression profiles likely differ across cell types in the analysed tissue.

**Fig. 4:**
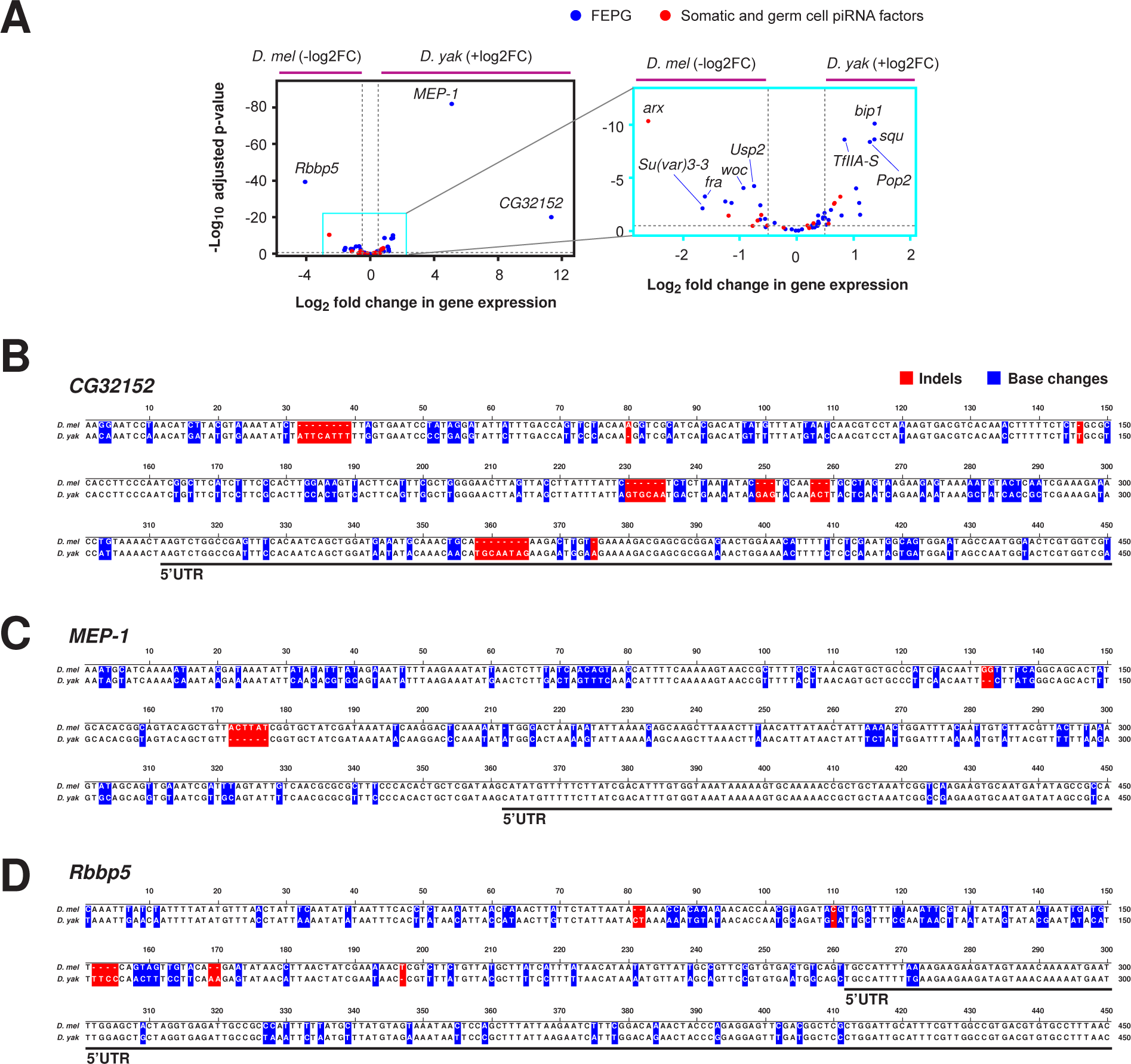
Analysis of differential gene expression of FEPG genes in *D. melanogaster* and *D. yakuba* ovaries. A) Volcano plot showing differential gene expression (DESeq2) between *D. melanogaster* and *D. yakuba* ovaries (RNA-seq data; Supplementary Dataset 1) for FEPG (blue) and the somatic and germ cell piRNA factors (red). Horizontal dashed lines indicate significance thresholds (adjusted p≤0.05). B-D) Alignment of the promoter regions of *CG32152, MEP-1* and *Rbbp5* for *D. melanogaster* and *D. yakuba* with indels (red) and base changes (blue) indicated as boxes. The black lines indicate gene 5’UTRs.

### Germ cell-specific genes show rapid evolution of promoters associated with divergent gene expression between species

To investigate whether the rapid evolution of promoters in piRNA factors expressed in germ cells is a common feature of germline-expressed genes, we used publicly available RNA-seq data from FACS-sorted vasa-GFP ovaries to identify genes specifically expressed in either germ cells or somatic cells of the ovary. First, using differential RNA-seq analysis of FACS-sorted vasa-GFP+ (germline) and vasa-GFP-(somatic) cells from cultured *D.mel* ovary GSC line (vasa-GFP;hs-bam;bamΔ86) (GSE119862) [41], we defined germline-enriched (vasa-GFP+; log2FC>0, p-val<0.01) and soma-enriched (vasa-GFP-; log2FC<0, p-val<0.01) genes (Fig. 5A and B). Next, using RNA-seq data from ovarian somatic cells (OSCs) (GSE160860) [42], we further filtered the soma-enriched genes that are highly expressed in OSCs. Additionally, we further filtered the germline-enriched genes that show expression in early fly embryos (0-2hr, modENCODE). In this way, we generated a stringent list of 107 genes specifically expressed in germ cells and of 59 soma-expressed genes (Supplementary Dataset 1).

**Fig. 5:**
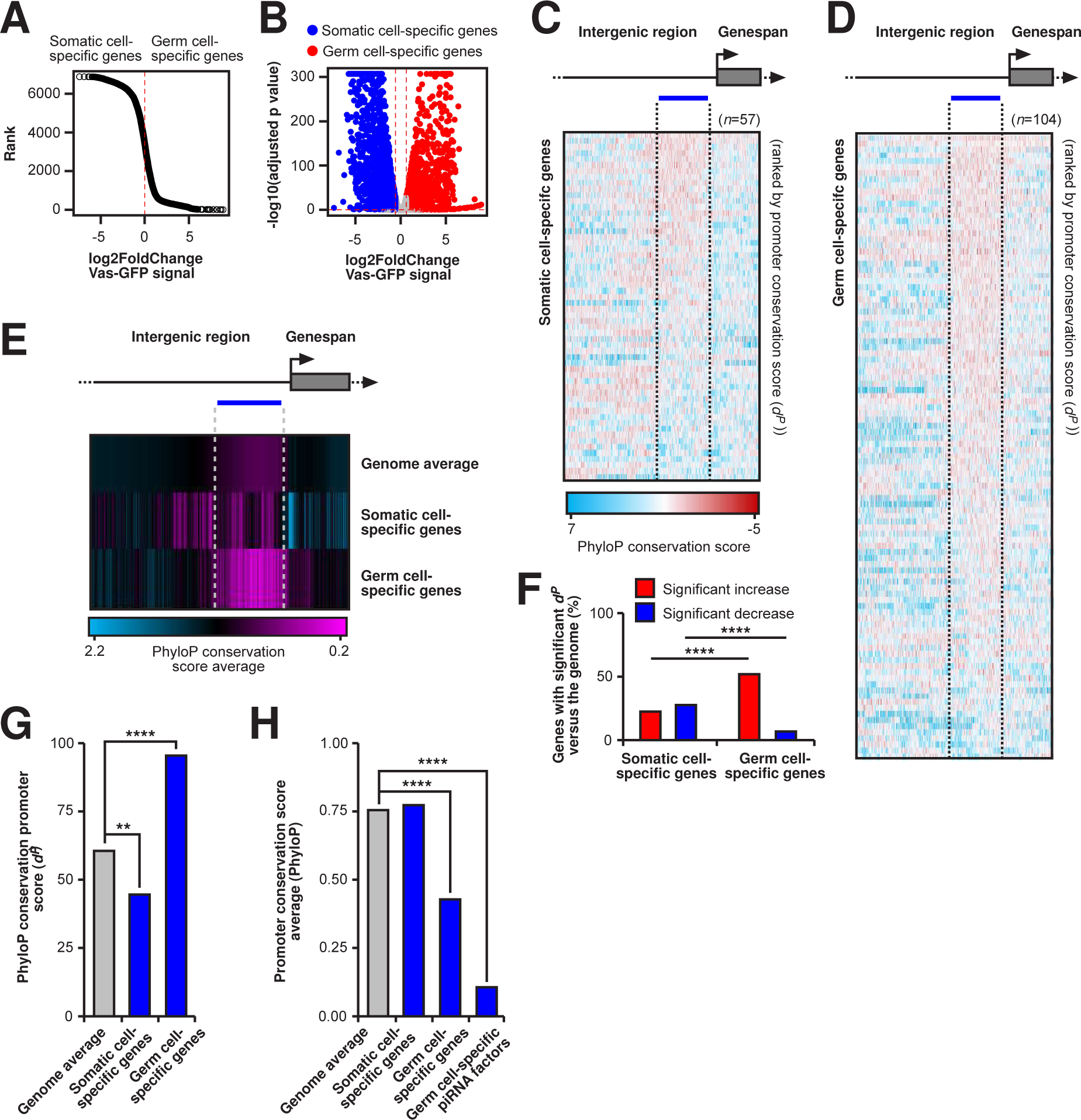
Germ cell-specifically expressed, but not somatic cell-specifically expressed gene promoters evolve rapidly and are accompanied by divergence in gene expression. A) Ranking of genes using vasa-GFP signal from FACS-sorted vasa-GFP+/- cells of cultured ovary GSC line (vasa-GFP;hs-bam;bamΔ86) (GSE119862) that defines germline-enriched (vasa-GFP+) and soma-enriched (vasa-GFP-) genes. B) Volcano plot showing differential gene expression between the FACS-sorted vasa-GFP+ (germline) and vasa-GFP-(somatic) cells from cultured ovary GSC line (vasa-GFP;hs-bam;bamΔ86) (GSE119862). C and D) PhyloP27way nucleotide scores for somatic cell-specific (C) and germ cell-specific (D) expressed genes ordered by promoter conservation *d^P^* scores. A region of 1000 nucleotides upstream and 300 nucleotides downstream are shown. Red represents lower conservation while blue represents higher. E) Average PhyloP27way nucleotide scores for all genes in the *Drosophila* genome, somatic cell and germ cell-specific genes. Purple represents lower conservation while blue represents higher. F) The percentage of somatic cell and germ cell-specific genes with significant promoter nucleotide changes (*d^P^*) versus all genes in the *Drosophila* genome average, divided by significantly increased (low conservation) or decreased (high conservation). Statistically significant differences from non-parametric chi-squared tests are indicated by asterisks (**** p≤0.0001 following Bonferroni correction). G and H) PhyloP27way conservation score averages (G) and PhyloP27way conservation promoter scores (*d^P^*) (H) for the 350 nucleotide promoter regions compared for all genes in the *Drosophila* genome, somatic cell-specifically expressed and germ cell-specifically expressed genes. Statistically significant differences from unpaired student t-tests (G) and from non-parametric chi-squared tests (H) are indicated by asterisks (** p≤0.01, **** p≤0.0001 following Bonferroni correction).

We then analysed sequence change accumulation in promoters using PhyloP data and calculated *d^P^* for all *D. melanogaster* genes and genes expressed specifically in somatic cells or the germline (Fig. 5C and D) or the average (Fig 5E). Germ cell-specific expressed genes significantly accumulated changes (*d^P^*) in their promoters compared to soma-expressed genes (Fig. 5F). Comparison of the average promoter PhyloP scores and *d^P^* conservation scores between the genome average and somatic or germ cell-specifically expressed genes equally revealed a significant increase in germ cell gene promoter nucleotide changes compared to the genome average (Fig. 5G and H).

Since rapid promoter evolution of germline piRNA silencing factors correlated with gene expression divergence in our previous analysis, we hypothesized that this was a general feature of germ cell genes. To test this, we compared gene expression divergence between somatic and germ cell-specific genes using publicly available RNA-seq data of *D. melanogaster* and *D. yakuba* ovaries (Fig. 6A) [38–40]. Notably, genes specifically expressed in germ cells showed slightly higher divergence in expression compared to soma-expressed genes (Fig. 6A). When *d^P^* conservation scores were plotted against the gene expression change between *D. melanogaster* and *D. yakuba*, the germ cell-specifically expressed genes showed slightly higher divergence in expression especially in the higher *d^P^* score groups (>50-75, >75-100) compared to soma-expressed genes, which showed relatively equal expression divergence across the different gene groups (Fig. 6C-E).

**Fig. 6:**
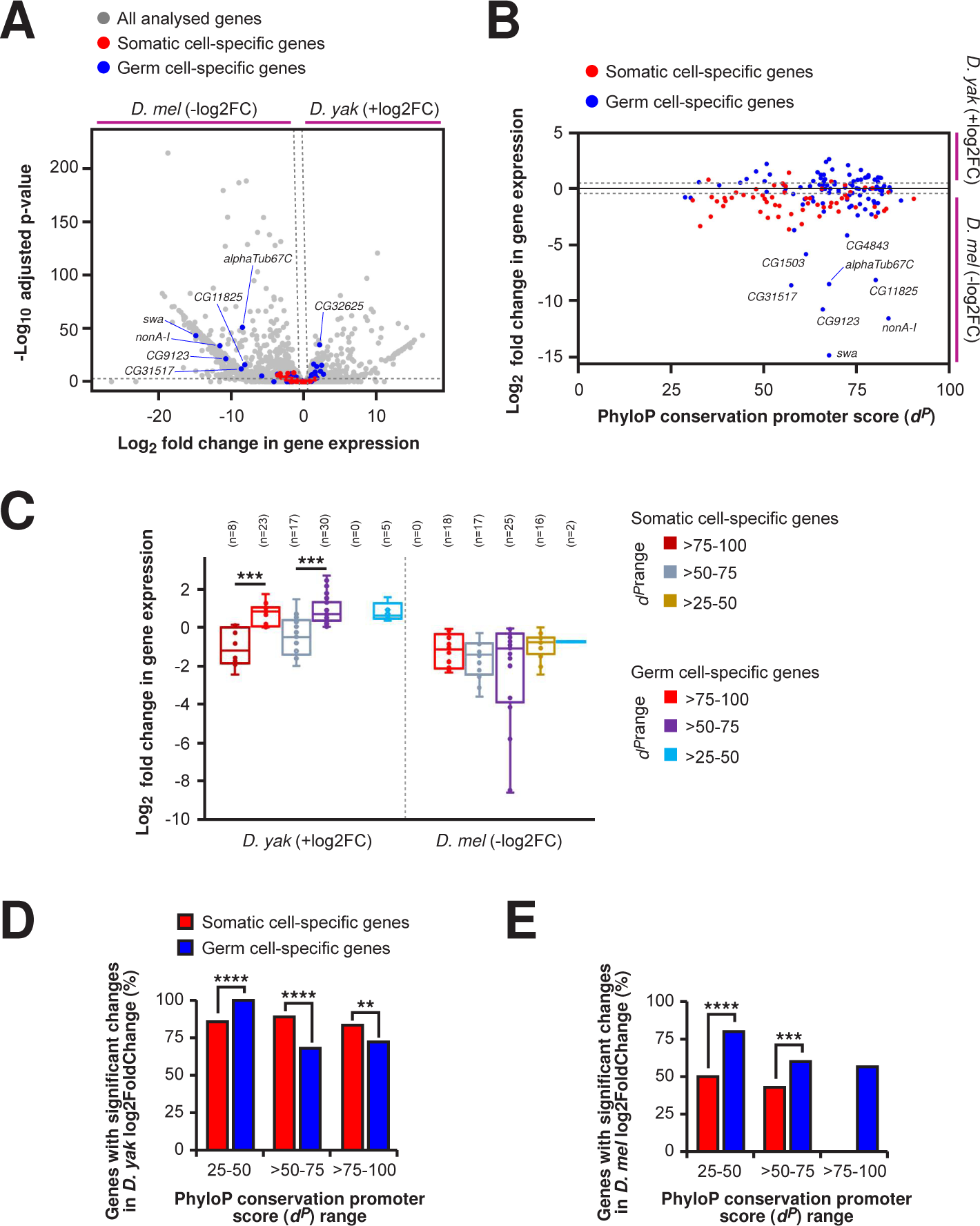
Analysis of gene expression divergence between germ cell and somatic cell-specifically expressed genes. A) Volcano plot of differential gene expression (DESeq2) between *D. melanogaster* and *D. yakuba* ovaries (RNA-seq data; Supplementary Dataset 1) for all (grey), germ cell-specific (blue) and somatic cell-specific (red) gene groups. Horizontal dashed lines indicate significance (adjusted p≤0.05). B) Scatterplot of individual gene *d^P^* scores plotted against fold-change expression differences between *D. melanogaster* and *D. yakuba* ovaries (RNA-seq data; Supplementary Dataset 1) for germ cell-specific (blue) and somatic cell-specific (red) gene groups. The grey dotted lines indicate log2Fold-change threshold values of ≥0.5 and ≤0.5. C and E) Comparison of the expression distribution and percentage of genes with significant changes in *D. melanogaster* log2FoldChange (≥0.5, D) or significant changes in *D. yakuba* log2FoldChange (≤0.5, E) expression changes in PhyloP conservation promoter score (*d^P^*) ranges 25-50, >50-75, and >75-100. Expression changes for germ cell-specific and somatic cell-specific gene groups were analysed between *D. melanogaster* and *D. yakuba*. Statistically significant differences from Mann Whitney U tests (C) and non-parametric chi-squared tests (D and E) are indicated by asterisks (** p≤0.01, *** p≤0.001, **** p≤0.0001 following Bonferroni correction).

## Discussion

Through genomic analysis of germline and somatic transposon silencing genes and their regulators we identify hot spots for rapid evolution in promoter regions (-30 to -380 nucleotides upstream of the TSS) of germ cell-specific (dual-strand clusters and the ping-pong cycle), but not soma-expressed piRNA pathway genes (cTGS and piRNA biogenesis). Further, we show that rapid promoter evolution is a general feature of germ cell-specific genes compared to those expressed only in the soma. Our analysis suggests that this mode of evolution in the germline pathway could be a general feature of at least insect evolution. Overall, our results point towards a key role for rapid evolution of gene promoters in the germ cell-specific piRNA pathway which could, coupled with other drivers of expression divergence, impact the expression of germline regulatory genes.

Transposon mobility has been attributed to the accumulation of sequence changes in promoter regions because of the presence of open chromatin around the transcription start site of genes [43]. Interestingly, in core NPC genes, rapid evolution mostly led to accumulation of indels [20, 21], while piRNA pathway genes display equal accumulation of indels and base changes. Likely, indels have more profound effects on changes in transcription than single nucleotide alterations. This feature might reflect that compromised transposon silencing causes sterility, hence not allowing for substantial changes in expression in piRNA processing genes. Likewise, stoichiometric changes in protein complexes can drastically alter complex functionality, e.g. the male-specific lethal (MSL) complex binds less to its targets when one component of the complex is missing [44, 45]. Hence, changes in the protein levels of individual piRNA factors likely have dominant effects on their capacity to silence transposon activity.

The occurrence of novel transposons can fundamentally impact species reproductive success through a phenomenon call hybrid dysgenesis [46]. Here, if a novel transposable element is transmitted by the male, female fertility is severely compromised and can result in sterility as a result of missing piRNA silencing of the novel invader. Essential to combat these novel active transposable elements is the ping-pong cycle, primed by transcripts from the novel transposon. Accordingly, females can be become resistant to such novel transposons over time through RNA transgenerational inheritance of piRNAs to build up ping-pong cycle amplification in germ cells [9, 10, 47].

Our examination of evolution in coding regions flagged germ cell-specific piRNA factors, as well as some pleiotropic piRNA processing associated complexes (EJC outer shell, EJC transient facts, NSL complex, SUMO modifiers, and Nuclear CBC) as under positive selection. This is unsurprising in the case of primary piRNA processing factors in the germline, as there are many examples of these factors being under selection [48–56]. For instance, replacement of *D. melanogaster rhino* and *deadlock* genes with *D. simulans* homologues results in non-functionality [49]. Here, *D. simulans* Rhino binding domains no longer bind to *D. melanogaster* Deadlock, resulting in failed localisation to piRNA clusters [49]. Such effects resulting from few amino acid changes act as powerful driving forces in diverging piRNA processing. In this setting, rapid promoter evolution of piRNA silencing genes could act as an additional factor driving changes in expression levels and factor stoichiometry for piRNA processing divergence, which may be difficult to explore when coupled with functional divergence. Intuitively, one would associate rapid promoter revolution with changes in expression, however, we observed minimal effects on expression. Since transposon silencing is essential, changes in cis regulatory elements of these regulatory factors is likely constrained by compensatory mechanisms and could be accompanied by changes in enhancer regions. Whether this is the case would required a detailed knowledge of relevant enhancers and transcription factor binding.

The pattern of changes observed in fast evolving promoters like germline specific piRNA factors and Nups resembles the outcome of a *P*-element mutagenesis experiment in the *egghead* (*egh*) locus coding for a glycosphingolipid biosynthesis enzyme [57]. Here, multiple base changes were observed in the region of the first promoter after *P*-element mutagenesis, but not an actual *P*-element insert, resulting in a sex peptide insensitive allele *egh^cm^*. Mutations in the *egh* gene result in pleiotropic phenotypes and the *egh^cm^* allele disrupts neuronal wiring required for the female post-mating response and development of the optic lobe [22, 57, 58].

Since establishing transposon resistance involves forced selection, genes whose expression changes result in pleiotropic effect, like channel Nup54, could combine adaptations in piRNA processing with changes in neuronal wiring resulting in altered behavioural preferences [20]. Of note, in *D. simulans*, a close relative of *D. melanogaster*, projections of *fruitless* P1 sensory neurons that control courtship have changed and alter mate preference [59].

The severe impact of hybrid dysgenesis on fertility likely limits fast evolution to species with high numbers of progeny. Perhaps this can explain the differences seen when comparing insect and mammalian promoters that are generally more conserved [26]. However, rapid promoter evolution has also been observed in primate promoters, suggesting a common mode of evolution that has yet to be explored [20, 21, 26, 60].

## Materials and methods

### Sequence/data retrieval and alignment

Gene and promotor sequences were retrieved from UCSC Genome Browser using the UCSC Table Browser sequence retrieval tool [33, 34]. A standardised region of 2000 bases upstream of the annotated gene TSS was exported for each gene to ensure inclusion of promoter regions. Sequences were imported and aligned with clustalW in MEGA11 [61]. PhyloP27way data were sourced from UCSC genome browser (genome.ucsc.edu) through the Table Browser tool [33, 34]. Data points were collected for a region of 1000 nucleotides upstream and 300 nucleotides downstream for each of the analysed genes. Genes without full species coverage when analysing evolution between the 5 *Drosophila* species, where ≥1 species had no conserved genomic sequence were removed from the analysis. Genes without full PhyloP coverage were removed from the analysis (where ≥1 nucleotide of the analysed genomic region contained the *D. melanogaster* sequence only).

### Molecular evolution of open reading frames analysis

Analysis of adaptive protein evolution was performed via MKTs [32] using the PopFly online database (imkt.uab.cat) which uses Drosophila Genome Nexus project sequence data [62–64]. Polymorphism and divergence data were collected from the ancestral Congo population, chosen since it is a sub-Saharan population with comparably high ancestral stability compared to other populations. The ‘Standard MKT’ test was used to calculate results for individual genes. Fisher’s Exact Test with significance defined as Bonferroni corrected p≤0.05 was used to calculate significance between nonsynonymous to synonymous polymorphisms within *D. melanogaster* or between *D. melanogaster* and *D. simulans*.

### Identification of promotor hot spots and comparison of substitution rates

All gene models were manually inspected in FlyBase using the JBrowse browser and dominant transcripts were chosen based on comparative modENCODE expression data [65]. Gene and promoter sequence alignments were trimmed to -1000/+300 nucleotides from the TSS of each gene. To calculate the frequency of sequence change hotspots in promoter sequences, we calculated the hotspot accumulation score *d* using alignments generated as described, or PhyloP data. Alignments were translated into ‘events’ for each species compared to *D. melanogaster*, where for each nucleotide in the sequence, 0 signified a conserved sequence and 1 signified a sequence change (base change or indel event) [66]. Events were calculated for all changes, as well as base changes and indels individually. The sum of events was calculated for concatenated gene groups and individual genes at each nucleotide position. A sliding event (*Se*) score was calculated from this using a sliding window of five bases along the sequence, from which heatmaps were generated. To calculate the percentage of events greater than the average control promoter *Se* score (*d*), a 350-nucleotide region upstream of the estimated TATA box region was analysed where the total number of *Se* scores exceeding the control group average sliding event score (*Se^C^*) was divided by the total number of events in that region (*N*).

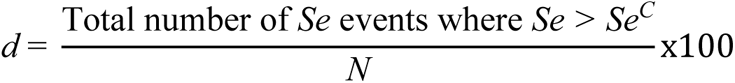

To calculate the promoter region *d* scores from PhyloP data, the total number of PhyloP (*p*) scores in the 350 nucleotide regions less than the control group average promoter region (*p^C^*) took the place of the total number of *Se* events where *Se* is greater than *Se^C^* (see equation below).

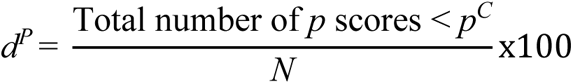

Significance was calculated using non-parametric chi-squared tests compared to the control group *d* score. Significance values where p≤0.05 with Bonferroni correction were considered statistically significant.

### Identification of somatic and germ cell-specifically expressed genes in *D. melanogaster*

We defined somatic and germline genes using several steps. First, using processed RNA-seq data from vasa-GFP+/- cells sorted from cultured ovary GSC line (vasa-GFP;hs-bam;bamΔ86) (GSE119862) [41], we defined germline-enriched (vasa-GFP+) using logFC>0, p-value<0.01 parameters and soma-enriched (vasa-GFP-) genes using logFC<0, p-value<0.01 parameters (Figures 4a and 4b). Next, using processed RNA-seq data from fly ovaries (modENCODE), we filtered genes that are expressed in adult fly ovaries using RPKM>10 parameter for both soma and germline genes. To stringently define the germline genes, we additionally used processed RNA-seq data from early fly embryos (0-2hr, modENCODE; mRNA-seq fly embryo 0-2hr; RPKM; mE_mRNA_em0-2hr; FBlc0000086) to further filter the germline-enriched genes using >9 RPKM parameter. To stringently define the somatic-specific genes, we additionally used processed RNA-seq data from ovarian somatic cells (OSCs) (n=4, GSE160860) [42] and filtered the soma-enriched genes using >20 RPKM to select genes that are expressed in OSCs as well as parameter <2 RPKM for early fly embryo (0-2hr, modENCODE) to filter out genes that are expressed in early embryos. This analysis generated a list of 107 germ cell-specifically expressed genes and of 59 somatic cell-specifically expressed genes (Fig. 4A and B, Supplementary Dataset 1).

### Comparative gene expression analysis between *D. melanogaster* and *D. yakuba*

Using publicly available raw RNA-seq data from *D. melanogaster* (n=3) and *D. yakuba* (n=4) ovaries, we trimmed adapter sequences from the raw reads using Cutadapt tool (v1.18, default parameters) and aligned the trimmed reads to the genome assemblies (dm6 and droYak2) using the RNA-seq aligner STAR (v2.7.3a). Gene counts were quantified using the featureCounts tool (Subread package v1.5.3). Orthologues genes were identified using blastDm2FB in UCSC Table Browser. Gene count normalization and differential gene expression analysis between the species were performed using DESeq2 (v1.30.1) with an additional species-specific gene lengths normalization step.

## Supplementary information

Supplementary materials include Fig. S1-4 and Supplementary Dataset 1.

## Authors’ contributions

DWJM and MS conceived and directed the project. DWJM performed genomic and evolutionary analysis. AA and DWJM analysed differential gene expression. DWJM wrote the initial draft and all authors contributed to the final manuscript. All authors read and approved the final manuscript.

## Acknowledgements

We thank the Soller lab for helpful discussion.

## Funding

This work was supported by the Medical Research Council to DWJM [MR/N013913/1] and the Biotechnology and Biological Science Research Council to MS. Work by BCN and AA was funded by Cancer Research UK [A21143] and the Wellcome Trust [110161/Z/15/Z].

## Availability of data and materials

All data generated or analysed during this study are included in the supplementary information files.

## Ethics approval and consent to participate

Not applicable.

## Consent for publication

Not applicable.

## Competing interests

The authors declare no competing interests.

## Figure Legends

**Supplementary Fig. 1:**
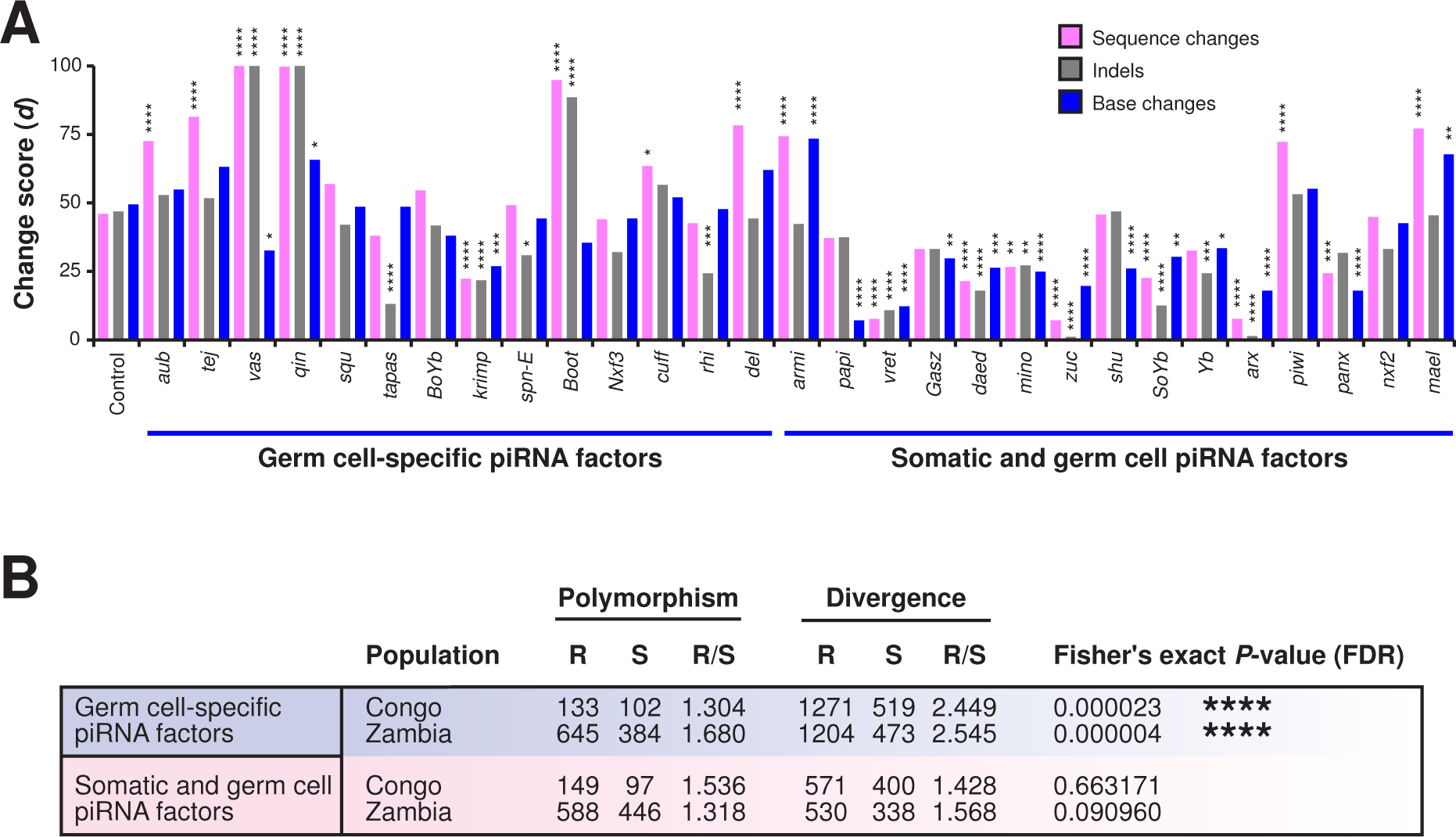
Germ cell-specific piRNA factors are under positive selection and accumulate rapid sequence changes in their promoters. A) Individual gene sequence change, indel, and base change *d* scores and statistically significant accumulation of sequence changes for the somatic piRNA factors and germ cell-specific piRNA factors. Statistically significant differences from non-parametric chi-squared tests are indicated by asterisks (* p≤0.05, ** p≤0.01, *** p≤0.001, **** p≤0.0001 following FDR correction). B) Evolution of coding regions analysed by MKT tests for polymorphisms and divergence between *D. melanogaster* and *D. simulans* for core genes in the somatic and germ cell transposon silencing pathways. R, replacement; S, synonymous. Statistically significant differences from Fisher’s Exact Tests are indicated by asterisks (**** p≤0.0001 following FDR correction).

**Supplementary Fig. 2:**
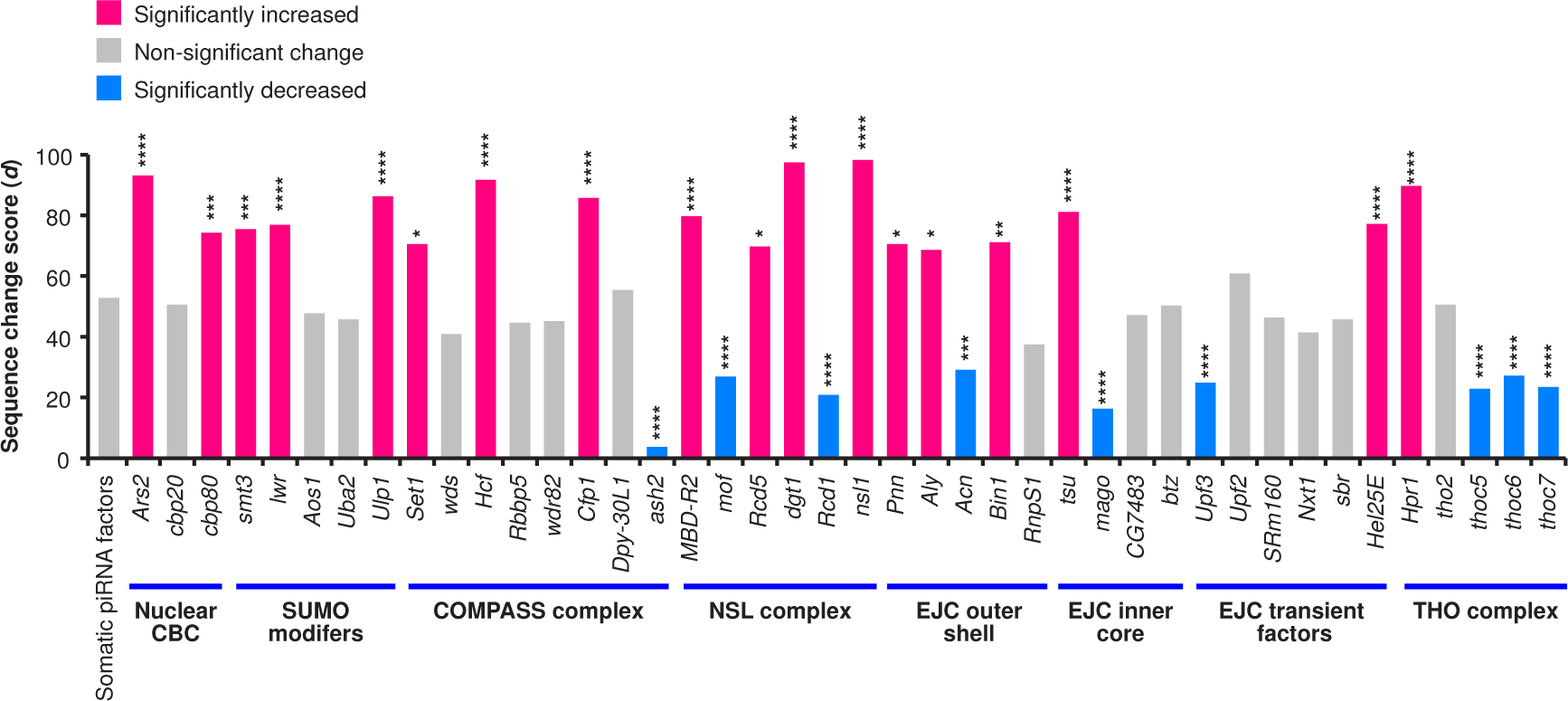
Genes coding for members of a subgroup of protein complexes implicated in piRNA silencing accumulate sequence changes in their promoters. Individual gene *d* scores and statistically significant accumulation of sequence changes for each gene separated by protein complex compared to the somatic piRNA factors. Statistically significant differences from non-parametric chi-squared tests are indicated by asterisks (* p≤0.05, ** p≤0.01, **** p≤0.0001 following Bonferroni correction).

**Supplementary Fig. 3:**
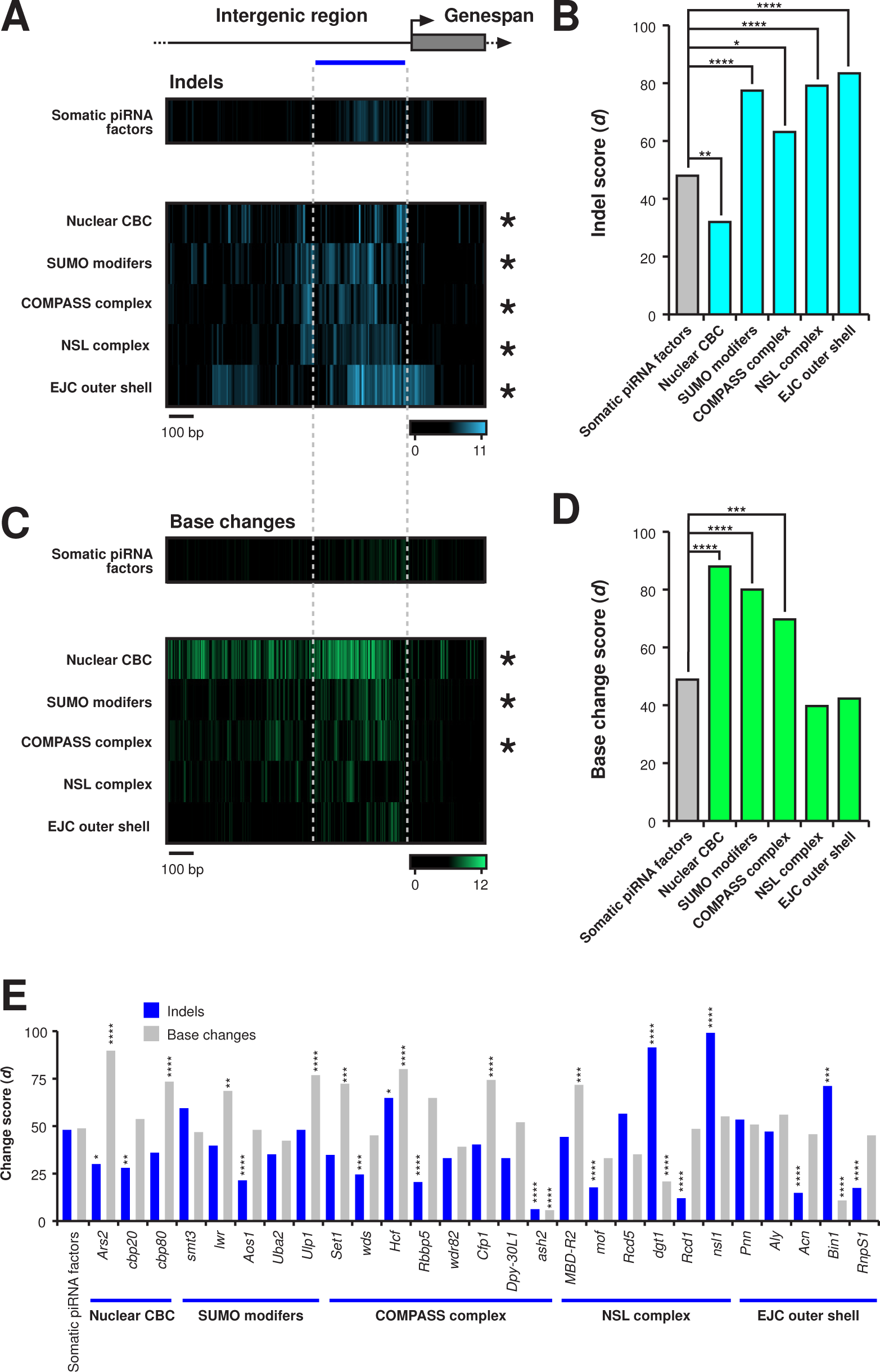
Promoter evolution of protein complex genes involved in germ cell transposon silencing. A and C) Heatmaps indicating indel (A, blue) or base change (C, green) accumulation in protein complex genes involved in germ cell transposon silencing compared to Yb body genes among closely related *D. melanogaster*, *D. simulans*, *D. sechellia*, *D. yakuba* and *D. erecta*. Regions of 1000 nucleotides upstream and 300 nucleotides downstream of the TSS were analysed. The blue line indicates the promoter region used for quantification of the substitution rate. B and D) Comparison of indel (B, blue) or base change (D, green) *d* scores for each of the protein complex gene groups compared to the somatic piRNA factors. Statistically significant differences from non-parametric chi-squared tests are indicated by asterisks (* p≤0.05, ** p≤0.01, *** p≤0.001, **** p≤0.0001 following Bonferroni correction). E) Individual gene *d* scores and statistically significant accumulation of indels or base changes for each gene separated by protein complex compared to the somatic piRNA factors. Comparison of indel (B, blue) or base change (D, green) *d* scores for each of the protein complex gene groups compared to the control group. Statistically significant differences from non-parametric chi-squared tests are indicated by asterisks (* p≤0.05, ** p≤0.01, *** p≤0.001, **** p≤0.0001 following Bonferroni correction).

**Supplementary Fig. 4:**
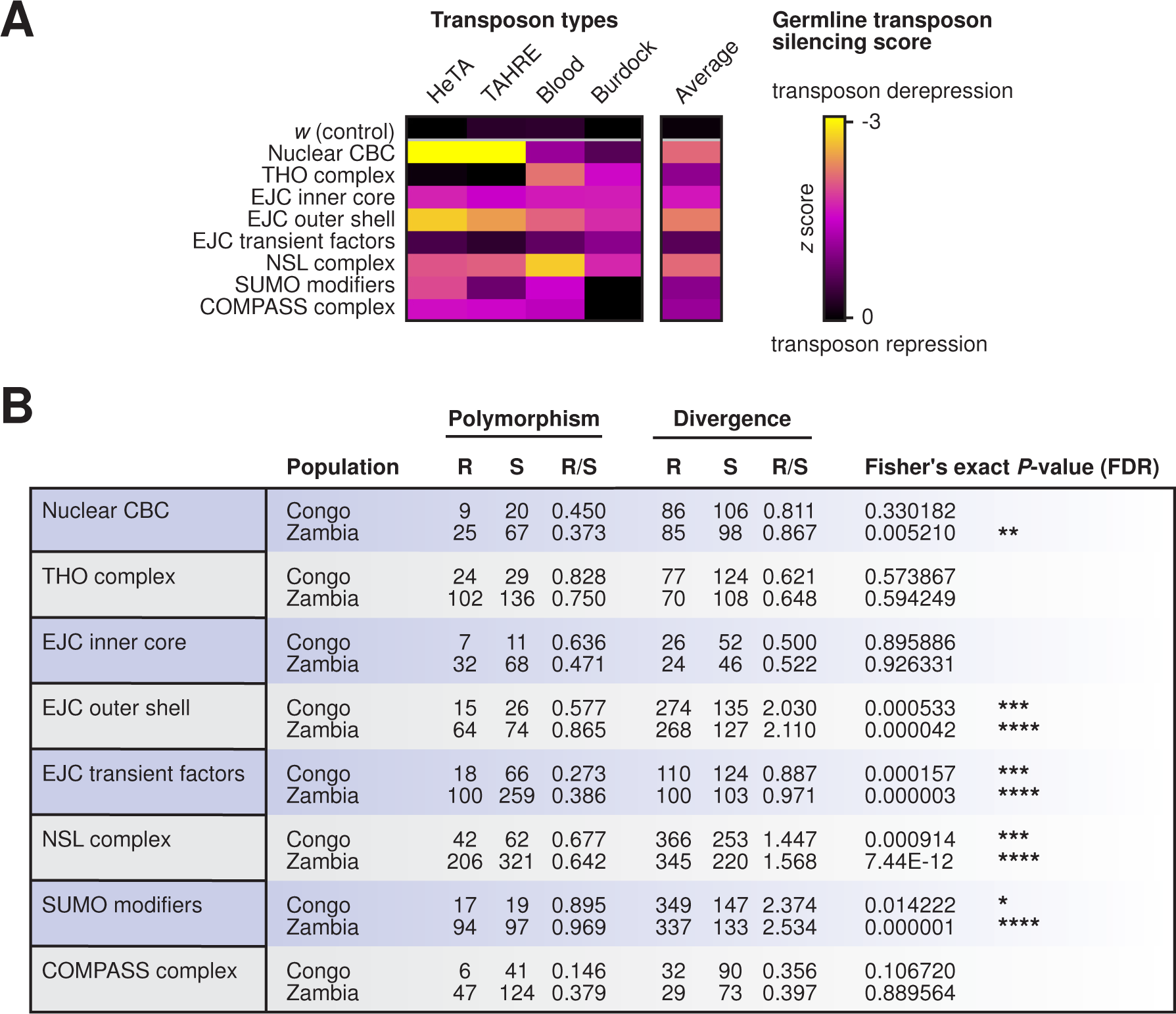
Coding sequences evolution of protein complex genes involved in germ cell transposon silencing. A) Impact of protein complex genes involved in germ cell transposon silencing on transposon de-repression. Unprocessed data were taken from [1]. Derepression scores (*z*) were concatenated for protein complexes in each transposon type (HeTA, TAHRE, blood, burdock) and an average of all. B) Evolution of coding regions analysed by MKT tests for polymorphisms and divergence between *D. melanogaster* and *D. simulans* for individual genes from the selected germline transposon silencing implicated protein complexes. R, replacement; S, synonymous. Statistically significant differences from Fisher’s Exact Tests are indicated by asterisks (* p≤0.05, ** p≤0.01, *** p≤0.001, **** p≤0.0001 following FDR correction).

## Notes

### Competing Interest Statement

The authors have declared no competing interest.

### Summary of Updates

Revised text and additional evolutionary and expression analysis.

